# Cryoelectron tomography of HIV-1 cell-cell transmission conjugates reveals a secluded environment for viral assembly and transfer

**DOI:** 10.64898/2026.06.11.725753

**Authors:** Jacob Jensen, Marcela Cunha, Broti Adhikary, Zaida Rodriguez, Iszavier Moe, Jasmine Beldin, Dmitry Lyumkis, Luiza Mendonça

## Abstract

Cell-cell transmission of HIV-1 (CCT) is a highly efficient mode of intrahost transmission that provides viral particles broad protection from antiretroviral agents including antiretroviral drugs, host restriction factors, and broadly neutralizing antibodies. Interestingly, the exact factors that grant viral particles increased protection and efficiency during intrahost spread remain unclear. Using a T cell co-culturing system and correlative cryoelectron tomography (cryoET) workflow, we investigated the architecture of HIV-1 CCT sites in a near-native, *in situ* context to address why CCT is efficient and grants this resistance. Contrary to previous models which suggest large, virus-packed sites, our 3D reconstructions reveal that CCT occurs within small intercellular spaces containing on average 1–5 viral particles. Surrounding these spaces are long, tight membrane interfaces that seclude the viral particles from the remaining extracellular environment. These observations suggest a model where spatial isolation may limit accessibility of antiretroviral agents to the virus-containing spaces thereby conferring protection to viral particles during transmission. To functionally validate these structural insights, we developed a quantitative flow cytometry strategy that decouples CCT conjugate formation from successful CCT infection through the analysis of multicell events. This methodology allows us to systematically compare the impact different molecules such as Env and CD4 have on HIV-1 CCT in a quantitative and population-level manner. Using this approach, we determined that initial HIV-1 CCT conjugate formation is partially dependent on CD4, but the Env involved in conjugate formation may differ from Env used for viral entry. Overall, these findings may prompt a rethinking of intervention strategies for HIV-1 CCT and have further implications with cytoplasmic trafficking, nuclear import, and host restriction factors.

## Introduction

HIV-1 can infect cells through two mechanisms. In free-virus transmission (FVT), a virus released from an infected cell diffuses until it finds, interacts and infects a susceptible target cell. With cell-cell transmission (CCT), the virus subverts the infected cell to turn it into a vector to find, interact and transmit viruses to susceptible cells with limited exposure to the intercellular space (1–5). This mode of transmission is considered one to two orders of magnitude more efficient than FVT (6–8), although the precise mechanism is still under debate. Viruses transmitted through CCT are resistant to some host restriction factors (9, 10), antiretroviral drugs (11–15) and broadly neutralizing antibodies (bnAbs) (1, 16–21), the latter with serious impact for current anti-HIV bnAbs-based therapy efforts (22).

Cell-cell transmission has been shown to be dependent on both viral and host factors (4, 7, 23, 24). One of the most important factors for successful CCT is the presence of the viral Envelope glycoprotein (Env) (25). It has been shown that cell-associated Env interacts with CD4 on the surface of target uninfected cells and is one of the factors that provides the intimate contact necessary for successful cell-cell transmission. Host cell integrins such as LFA-1, ICAM-1 and ICAM-3 have also been shown to be involved in CCT (7) but it remains unclear whether their interaction is crucial for stable intimate contact (26), polarization of proteins to the cell-cell interface (7), or both.

Previous investigations of HIV-1 CCT have utilized resin-embedded electron microscopy, fluorescent imaging, and flow cytometry to understand the mechanism of the increased efficiency of CCT (5, 27–31). These data indicate that CCT involves the polarization of Gag and Env to the cell-cell interface, suggesting that a large number of viruses might be transmitted at once (12, 16). This high local concentration of viruses could explain why drugs, restriction factors and bnAbs fail to block transmission, as there may be viruses that are not efficiently neutralized and escape to achieve successful transmission and infection of the target cell.

Electron microscopy of resin-embedded samples has shown that the cell-cell transmission sites are created by interdigitation of the plasma membranes of infected and uninfected cells, creating large compartments of scalloped membranes containing dozens of viral particles per site (5, 23).

Infection efficiency of CCT has been mostly quantified using cytometry assays to measure the number of newly infected cells after 24 hours of co-culturing infected cells with uninfected cells. Blocking agents have also been incorporated in these cytometry assays to identify one or more interactions that are needed for CCT (32–34). However, these assays cannot assess whether an interaction is required for CCT conjugate formation between infected and target cell or if the interaction is necessary for successful infection of the target cell, or both.

Novel advances in cryoelectron microscopy (cryoEM) have enabled the visualization of host-pathogen interactions at sub-nanometer resolution in a near-native, hydrated condition (35–37). However, the application of cryoEM, specifically cryoelectron tomography (cryoET), to CCT has been historically hindered because the cellular interfaces, where transmission occurs, are far too thick for direct electron penetration. To overcome this, recent correlative imaging workflows have integrated focused ion beam milling under cryogenic conditions (cryoFIB/SEM) (36, 38, 39). This technique allows for the precise thinning of cellular samples into transparent ‘lamella’ of less than 200 nm in thickness. By combining fluorescent targeting with cryoFIB/SEM, it is now possible to perform accurate targeted imaging and navigate the low probability of capturing transient, stochastic events within the small volume of a vitrified sample (40, 41). Altogether, these advancements have made it possible to localize the membrane contacts between HIV-1 infected and uninfected cells and resolve the macromolecular architecture of CCT sites without the artifacts associated with traditional resin-embedding microscopy.

In this study, we present a correlative cryoET imaging workflow to image CCT sites in near-native, hydrated conditions and characterize their ultrastructure. We also utilized a novel flow cytometry strategy to understand the importance of the Env-CD4 interaction on initial CCT conjugate formation, as well as CCT infection. Our *in situ* cryoET imaging showed small and secluded intercellular spaces that harbor 1-5 viral particles in various assembly and maturation states. These CCT conjugates, but not subsequent infection, were resistant to Env blockade but not resistant to cellular CD4 blockade or Env knockout.

## Materials and Methods

### Cell Culture

HEK 293T/17 and TZM-bl cells were maintained in DMEM high glucose (Gibco #11995073) supplemented with 10% fetal bovine serum (FBS) (Gibco, #A5256801) at 37°C and 5% CO_2_. SupT1 R5 cells and primary CD4+ T lymphocytes were maintained in RPMI-1640 media (Gibco, #A1049101) supplemented with 10% of FBS at 37°C and 5% CO_2_. HEK 293T/17 cells were obtained from ATCC (#ACS4500). Sup T1 R5 cells were a kind gift from Dr. Eric Freed’s lab. TZM-bl cells were obtained through BEI Resources, NIAID, NIH (#HRP-8129).

### Plasmids and Molecular Cloning

HIV-1 infectious molecular clone NL4-3 IRES-eGFP (#ARP-11349, NL4-3 IRES-eGFP Infectious Molecular Clone pBR43IeG-nef+) and NL4-3 (#ARP-114, Strain NL4-3 Infectious Molecular Clone pNL4-3) were obtained through the NIH HIV Reagent Program, Division of AIDS, NIAID, NIH, contributed by Dr. Jan Münch, Dr. Michael Schindler and Dr. Frank Kirchhoff and by Dr. M. Martin, respectively (42, 43). **Δ**Env IRES-eGFP is an Envelope-deleted, non-infectious version of NL4-3 IRES-eGFP generated by restriction digestion with PsiI-v2 (New England Biolabs, #R0744S) followed by re-ligation. VSV-G envelope expressing vector pMD2.G was a gift from Didier Trono (Addgene plasmid #12259).

### Virus Production and Titration

HEK 293T/17 were seeded in 75 cm^2^ flasks, transfected with 10 μg of NL4-3 IRES-eGFP plasmid or transfected with 10 μg of **Δ**Env IRES-eGFP plasmid with 2 μg of pMDG.2, using GenJet Transfection Reagent (Signagen Laboratories, #SL100489) after reaching 85% confluency. For immunofluorescence assays, 10 μg of NL4-3 plasmid was used for HIV-1 viral particle generation. Supernatants containing viruses were collected 48 hours post-transfection, filtered and viral particles were concentrated using Lenti-X concentrator (Takara Bio, USA, # 631231) following manufacturer’s instructions. Viral aliquots were stored at -70°C until needed. TZM-bl cells were plated 24 hours prior to titration at 4×10^4^ cells per well on a 96-well plate. Viral stocks were then serially diluted in DMEM with 2% FBS and the plate was incubated for 48 hours at 37°C and 5% CO_2_. Following incubation, cells were washed with PBS, fixed with 2% paraformaldehyde (PFA) solution, and then stained using X-gal staining solution for 24 hours at 37°C, as previously described (44). The total number of blue foci were counted in the most diluted well to have at least 30 blue foci and the infectivity of viral aliquots was determined (infectious particles/ml) for later experiments.

### Isolation of human CD4+ T lymphocytes from peripheral blood mononuclear cells (PBMCs)

Human peripheral blood mononuclear cells (PBMCs) were obtained from Trimacones (Innovative Blood Resources) from healthy donors. The gender distribution of the samples was 50% female and 50% male. PBMCs were isolated using Histopaque-1077 density gradient (Sigma-Aldrich, #10771) by centrifugation at room temperature, 950 xg for 20 min. CD4+ T cells were isolated from PBMCs by negative magnetic selection using microbeads and the MidiMACs separator (MiltenyiBiotec, Germany), according to the manufacturer’s instructions. After separation, cells were kept in culture in RPMI media containing 10% FBS and stimulated with 2.5 μg/ml phytohemagglutinin (PHA) (ThermoFisher, #00-4977-03) and 25 ng/ml interleukin 2 (IL-2) (Sigma-Aldrich, #H7041) for three to four days for the activation of CD4+ T lymphocytes. Cells were then used promptly in subsequent experiments.

### Cell-cell Transmission (CCT) Assay

6×10^5^ SupT1 R5 or primary CD4+ T lymphocytes were centrifuged in virus-containing unsupplemented RPMI-1640 (ThermoFisher #A1049101) media per condition at 1,200 xg for 2 hours at an MOI of 0.5-1 and 4, respectively. Post spin, an equal volume of RPMI-1640 media containing 20% FBS was added to obtain a final concentration of 10% FBS. After 24-48 hours of recovery at 37°C, infected cells were co-cultured with 3×10^5^ target SupT1 R5 or primary CD4+ T lymphocytes previously labelled with 1 μM CellTracker Deep Red (CTDR) (Invitrogen, #C34565) according to the manufacturer’s instructions. If required, cell populations were treated with soluble CD4 (sCD4) or anti-CD4 antibody SIM.2 30 min prior to co-culturing. sCD4 was administered at 5 and 20 μg/ml to infected cells. SIM.2 was administered at 0.1, 1, and 10 μg/ml to target cells. Co-cultures were maintained with RPMI-1640 containing 5% FBS at 37°C and 5% CO_2_. After incubation for 0, 3 or 24 hours, co-cultures were fixed with a final concentration of 4% PFA for 30 minutes at room temperature and used promptly in downstream methods. sCD4 was obtained through the NIH HIV Reagent Program, Division of AIDS, NIAID, NIH: Human Soluble CD4 Protein, Recombinant from CHO Cells, (#ARP4615), contributed by Progenics Pharmaceuticals, Inc. The Anti-Human CD4 Monoclonal SIM.2 was obtained through the NIH HIV Reagent Program, Division of AIDS, NIAID, NIH, (#ARP-723), contributed by Dr. James E.K. Hildreth.

### Immunofluorescence Confocal Imaging

6×10^5^ NL4-3 infected SupT1 R5 cells and 3×10^5^ uninfected CTDR+ SupT1 R5 target cells were generated as previously described in CCT assay methods. After 3 hours, co-cultures were fixed with 4% PFA for 30 minutes at room temperature and permeabilized with saponin 0.1% (ThermoFisher, #J63209.AK) for 10 minutes, followed by a blocking step with 2% normal goat serum and 1% BSA for 15 minutes. The cells were incubated overnight with the broadly neutralizing anti-Env antibody 2G12 (BEI Resources, #ARP-1476) and with conjugated antibodies for detection of p24 (KC57-FITC Conjugate) (BEI Resources, #ARP-13450) and human CD4 (CD4-APC Conjugate) (Miltenyi #130-113-222) at 1:200 dilution in a solution of PBS, 0.05% saponin, 1% BSA and 2% normal goat serum. After three washing steps with PBS, cells were incubated with Alexa Fluor 532 conjugated anti-human secondary antibody for 2 hours (1:200 dilution), and Hoechst 33442 was added for 10 minutes (Stemcell Technologies #100-1540). The cells were washed three times and slides were mounted using mounting medium (Ibidi #500001) and 1.5 mm coverslips (EMS #72224-01). The cell-cell transmission sites were analyzed with a NIKON AX R confocal microscope with the NIS Elements software. Image adjustments and montage were performed in ImageJ Fiji v.1.54p (45).

### Transwell Assay

6×10^5^ NL4-3 IRES-eGFP infected SupT1 R5 cells and 3×10^5^ uninfected CTDR+ SupT1 R5 target cells were generated as previously described in CCT assay methods. Infected cells were seeded on a transwell membrane insert containing 0.4 μm membrane pores (Corning, #3470) to limit direct contact with the target cell population. After 24 hours, cells were fixed using 4% PFA at room temperature for 30 minutes. Samples were then read using the FACSymphony A3 cytometer with emphasis on the number of all GFP+CTDR+ events. Data was analyzed using FlowJo v10.10.0 (Waters Biosciences).

### CCT Sample Vitrification

Following 3-hour incubation of CCT events, 2-5 μl of co-culture sample was added to glow-discharged, carbon-coated Quantifoil G300F1 R2/2 EM gold grids (Quantifoil). Sample was added to the carbon surface of the grids, and 1 μl of PBS containing 10% glycerol was added to the gold surface. The environmental chamber was kept at 80% humidity and 25°C during blotting from the gold side and subsequently plunge-frozen using the Leica GP2 system (Leica Microsystems) into liquid ethane at -180°C. The grids were stored in liquid nitrogen before using in downstream imaging.

### Cryo Correlative Light and Electron Microscopy (cryoCLEM)

Vitrified samples were imaged using the Leica cryoCLEM widefield microscope. Fluorescence signals were captured using a 50x / 0.90 NA cryo-objective. A 3-channel (GFP, CTDR and Brightfield) full grid map was acquired using the Leica LAS X software. To compensate for grid tilt and maintain focus across the entire area, a manual focus map was generated prior to acquisition, defining a single optimal Z-height for each tile. Infected donor cells (GFP+) and target cells (CTDR+) were identified, and the coordinates of CCT conjugates were mapped relative to the G300F1 grid indexing visual landmarks. These manual registrations were used to relocate and guide site-specific cryoFIB/SEM milling of the identified interfaces.

### Cryo Focused Ion Beam Milling and Scanning Electron Microscopy (cryoFIB/SEM)

Vitrified cell-cell interface lamellae were prepared using a ThermoFisher Scientific Aquilos 2 dual-beam cryoFIB/SEM equipped with a cryotransfer system and rotatable cryostage cooled at −191 °C. Prior to milling, grids were coated with an organometallic platinum layer using the GIS for 5–6 sec. Cell-cell interfaces with at least one infected and target cell positioned near the centers of grid squares were manually selected for thinning, guided by the previously acquired fluorescence grid maps loaded into MAPS software. Thinning was conducted via AutoTEM 5 (Thermo Fisher Scientific) in a stepwise manner, with the ion beam (30 kV) current reduced from 0.5 nA to 30 pA. The final target thickness of the lamellae was 200 nm.

### Cryoelectron Tomography (cryoET)

Lamellae-containing grids were transferred to a FEI Titan Krios G2 or FEI Krios G4 (Thermo Fisher Scientific) electron microscope operated at 300 kV and equipped with a Gatan BioContinuum 20eV-wide energy filter and K3 detector (Gatan), SelectrisX 10eV-wide energy filter with a Falcon 4 detector (Thermo Fisher Scientific) or a SelectrisX 10eV-wide energy filter with a Falcon 4i detector (Thermo Fisher Scientific). A 100 µm objective aperture was inserted. Membrane-membrane interface regions were selected for acquisition. Tilt series were recorded using either SerialEM software (46) or Tomo5 (ThermoFisher) with a nominal magnification of 53k, 64k, or 81k and a physical pixel size of 1.68, 1.965 or 1.562Å/pixel, respectively. The target defocus value for all imaging was set from -3 to -5 µm. The pre-tilt of the lamellae was applied, and a dose-symmetric scheme was used for all tilt series, ranging from -60° to +60° with an increment of 3° and groupings of 3. A target of 41 projection images with 8-100 movie frames each were collected for each tilt series, with an exposure time of 0.2-0.7 seconds, resulting in a total dose of ∼120 e/Å^2^. Frames were saved in electron event representation (.eer) format (47) or in TIFF (.tiff) format with no gain normalization.

### CryoET Data Processing

Collected cryoET tilt series were corrected for beam-induced motion and split into even/odd frames using MotionCor2 v.1.6.4 (48). Resulting stacks were then aligned and tomogram reconstructions generated using AreTomo2 v.1.1.2 (49). Prior to denoising, CCT site dimensions and particle composition were determined using IMOD v. 5.1.3 (50). For denoising, cryoCARE v.0.3.0 (51) with default parameters was run and tomograms subsequently segmented using napari v.0.7.0 (52). Visualization of segmentation was done using ChimeraX v.1.11.1 (53).

### Resin-embedded Transmission Electron Microscopy

Fixed co-cultures of 6×10^5^ NL4-3 IRES-eGFP infected SupT1 R5 with 3×10^5^ target CTDR+ SupT1 R5 were washed three times with a 0.1M sodium cacodylate buffer at pH 7.2 with 5 minutes between spin downs. Buffer was replaced with a 2.5% glutaraldehyde in 0.1M cacodylate buffer at pH 7.2 and cells were left at room temperature for 40 minutes. Following incubation, cells were then washed again three times with a 0.1M cacodylate buffer at pH 7.2 with 5 minutes between washes. Cells were then treated with a 1% osmium tetroxide in 0.1M cacodylate buffer at pH 7.2 for 30 minutes at room temperature in the dark followed by another set of three washes with 0.1M cacodylate buffer at pH 7.3. Cells were then washed with 30%, 50%, and 70% ethanol for 5 minutes each, stained with uranyl acetate for 30 minutes in the dark, and then washed again with 80%, 95% and 100% ethanol, each step being washed twice for at least 5 minutes. Ethanol was replaced with propylene oxide followed by embedding in epoxy resin. Ultrathin sections of 70 nm were cut and collected on TEM grids. Sections were imaged using a FEI Tecnai Spirit Bio-Twin transmission electron microscope and TEM Imaging and Analysis software (ThermoFisher).

### Imaging Cytometry

6×10^5^ NL4-3 IRES-eGFP infected or 6×10^5^ **Δ**Env IRES-eGFP-infected SupT1 R5 cells were co-cultured with 3×10^5^ uninfected CTDR+ SupT1 R5 target cells as previously described in CCT assay methods. After 3 and 24 hours, co-cultures were fixed with 4% PFA for 30 min at room temperature and resuspended in 50uL of PBS. A target of 1,000 events was collected using the Cytek® Amnis® ImageStream®X Mk II Imaging Flow Cytometer. Images were taken using a 40x objective with 488 nm (20 mW) and 642 nm (0.1 mW) excitation lasers. Channel 1 was recorded for Brightfield (457/45 nm), Channel 2 for GFP (533/55 nm) and Channel 5 for CTDR (702/85 nm). Resulting images were analyzed using the IDEAS software v.6.2.187.0 (Amnis/EMDmillipore). To determine if cells were syncytia based on size, we first calculated the average diameter of a **Δ**Env IRES-eGFP-infected SupT1 R5 cell as well as the average diameter of an uninfected target CTDR+ cell. Using these diameters, we estimated the average volume of each cell and then added these volumes together to get a volume equivalent to one syncytia. From this syncytia volume, we then calculated the average diameter of a syncytia. For classifying, we used one standard deviation below the average syncytia diameter as a threshold where any diameters greater than the threshold were considered a syncytia. Error was propagated through each calculation accordingly.

### Free-virus Transmission (FVT) Assay

6×10^5^ SupT1 R5 or 6×10^5^ primary cells were centrifuged in virus-containing unsupplemented RPMI-1640 (ThermoFisher #A1049101) media per condition at 1,200 xg for 2 hours at an MOI of 1 and 4, respectively. Treatments with 5 and 10 μg/ml s0oluble CD4 (sCD4) or 0.1,1, and 10 μg/ml anti-CD4 SIM.2 antibody were administered right before centrifugation. Controls were treated with PBS. Post spin, an equal volume of RPMI-1640 media containing 20% FBS was added to obtain a final concentration of 10% FBS. After 24 hours of recovery at 37°C, cells were fixed with 4% PFA and quantified for GFP expression using the FACSymphony A3 cytometer. Results were analyzed using FlowJo version 10.10.0 (Waters Biosciences).

### Conventional Flow Cytometry

Flow cytometry was used for the evaluation of multicell conjugate formation in 3-hour co-cultures between infected cells (GFP+) and target cells (CTDR+), new GFP+CTDR+ single cell infections in co-cultures after 24 hours, or new GFP+ single cell infections resulting from FVT. All samples were gated to remove small non-cell/debris events and very large clusters. To achieve reliable separation between single and multicell events, FSC-W vs. FSC-A, SSC-W vs. SSC-A and FSC-W vs. SSC-W plots were created and single cells gated according to uninfected controls. Events were considered single cells if found within all three gates. All other events were considered multicell conjugates. For 3-hour co-cultures, multicell conjugates were assessed for the number of GFP+CTDR+ events. For 24-hour co-cultures, only single cells were assessed for the number of GFP+CTDR+ events. GFP+CTDR+ events for both 3 and 24-hour co-cultures were normalized to the initially added infection population. For FVT, only single cells were assessed for the number of GFP+ events. A FACSymphony A3 Cell analyzer and FACSDiva software v.9 was used for the data collection (Waters Biosciences). Analysis was performed in FlowJo software v10.10.0 (Waters Biosciences).

### Statistical Analysis and Data Plotting

Statistical analysis was performed using GraphPad Prism Software v.11. Results were presented as the mean and ± standard deviation (SD). Comparisons between two datasets were performed using a two-tailed Student’s *t*-test whereas comparisons among groups were analyzed using one-way ANOVA with Tukey’s multiple comparisons test. *p*-values less than 0.05 were considered statistically significant. Graphs were generated using Python v.3.14.2 and Python packages Matplotlib v.3.10.8 (54), NumPy v.1.26.4 (55), and pandas v.2.3.3 (56).

## Acknowledgements

We thank the Mansky lab at the University of Minnesota for providing access to the cryoCLEM and plunge freezing instrumentation. We thank Dr. Janet Iwasa and the Animation Lab at the University of Utah for partnering to create the proposed cell-cell transmission model animation. We also thank the University of Minnesota Cytometry Core Facility and the University of Wisconsin Carbone Cancer Center Flow Cytometry Laboratory (supported by P30 CA014520) for the use of their facilities and services. We gratefully acknowledge access to the Salk Institute Aquilos and Leica cryoCLEM systems, supported by startup funds to D.L. The authors acknowledge the facilities of the UC San Diego cryoEM facility, part of the Goeddel Family Technology Sandbox, along with the scientific and technical assistance of facility staff members Dr. Mariusz Matyszewski and Dr. Inga Kuschnerus. Parts of this work were carried out in the Characterization Facility, University of Minnesota, which receives partial support from the NSF through the MRSEC (DMR-2011401) and the NNCI (ECCS-2025124) programs. A portion of this research was supported by NIH grant U24 GM139168 and performed at the Midwest Center for CryoET (MCCET) and the CryoEM Research Center in the Department of Biochemistry at the University of Wisconsin-Madison. This work was primarily supported by the National Institute of Allergy and Infectious Diseases (NIAID) of the National Institutes of Health under award numbers 1R21AI191967-01A1, U54AI170752, and 1U54AI170855-01 and the Gilead HIV Research Scholars Program awarded to L.M. Z.R. was supported by the Pathways in Biological Sciences (PiBS) NIH T32 Training Grant (T32 GM133351) and the Mary K. Chapman Foundation. D.L. was supported by NIH/NIAID grants R01 AI150998, R01 AI178849, U01 AI136680, 1R21AI197945, R01 AI184419, NCI CCSG P30 CA014195, as well as the Margaret T. Morris Foundation and Hearst Foundations. J.J. was supported by the T32 AI83196 Training Grant from the Institute for Molecular Virology at the University of Minnesota.

## Results

### Co-culturing of T cells results in polarization of cellular and viral proteins

To first assess HIV-1 CCT, we needed to confirm that our co-culturing system was adequate for generating productive CCT events. To accomplish this, we co-cultured NL4-3 infected and target SupT1 R5 cells together for 3 hours at a 2:1 ratio and stained for Gag, Env, and CD4. These proteins have been previously described to be localized to the cell-cell interface during HIV-1 CCT (4, 9, 57). Using confocal fluorescence microscopy, we confirmed the polarization of CD4 in the target cells and Env in the infected donor cell to the cell-cell interface at 3 hours (Figure 1A). This localization of viral and cellular proteins confirms that our co-culturing system can successfully produce CCT sites by 3 hours of incubation.

**Figure 1.**
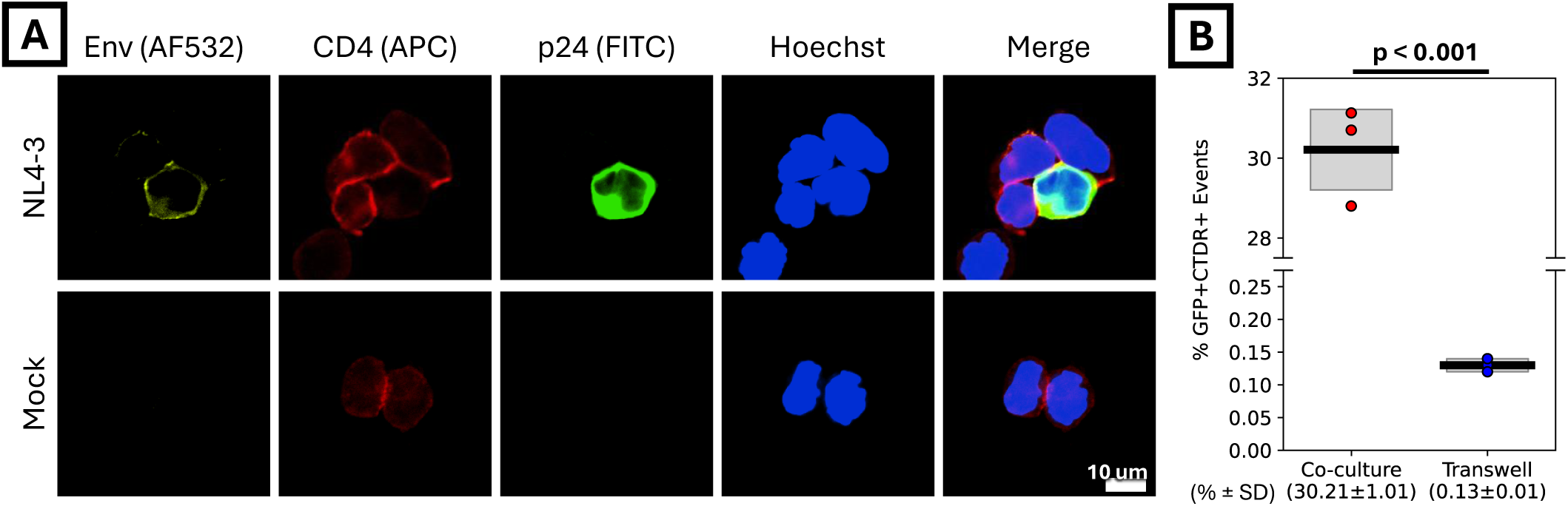
Immunofluorescence microscopy and co-culture validation of HIV-1 CCT. (**A**) SupT1 R5 donor cells infected with the HIV-1 molecular clone NL4-3 were co-cultured with uninfected SupT1 R5 target cells for 3 hours at 37°C. Cells were fixed, permeabilized, and immunostained for HIV-1 Env (yellow), CD4 (red), and p24/Gag (green), with nuclei counterstained using Hoechst 33342 (blue). (**B**) Percentage of newly infected target SupT1 R5 cells after 24 hours. NL4-3 IRES-eGFP infected SupT1 R5 cells (GFP+) were either co-cultured with CellTracker Deep Red stained target SupT1 R5 cells (CTDR+) or within a transwell system where infected and target cells were separated by a 0.4 µm membrane to block cell-cell contact. Data is representative of two biological replicates. Black bar represents the mean and the gray box represents standard deviation, which are also given below the x-axis. *p*-values calculated using two-tailed unpaired *t-*test.

### Free virus transmission is negligible in comparison to cell-cell transmission

To assess whether free-virus transmission (FVT) had any impact on infection in our co-culturing system, we conducted a transwell experiment separating the infected and healthy T cell populations from one another using a 0.4 μm membrane. SupT1 R5 T cells were infected with NL43 IRES-eGFP, viruses produced from a HIV-1 bicistronic clone that expresses soluble GFP using an Internal Ribosome Entry Site (IRES) downstream of intact *nef* (43). The IRES system downstream of the *nef* ORF ensures that there is minimal interference between GFP and other viral proteins during HIV-1 infection but still allows for identification of infected cells. The infected donor cell population was added to the upper section of the transwell insert. CellTracker Deep Red-stained SupT1 R5 cells were then added to the bottom of the transwell plate below the membrane. The plate was incubated for 24 hours and the CTDR-stained lower SupT1 R5 cells were assessed for GFP expression using flow cytometry. A parallel co-culture where cells were not separated by a membrane was also performed. By 24 hours, 30.21% of all events from the unimpeded co-culture were GFP and CTDR positive while in the CTDR stained cells separated by a transwell, we observed only 0.13% of events being newly infected (Figure 1B). This statistically significant difference indicates that FVT makes up a small fraction of successful infectious events in our co-cultures and that CCT is the dominant mode of transmission in our system.

### HIV-1 CCT occurs in small and secluded intercellular spaces containing few viral particles

We next applied a correlative cryoimaging workflow to image CCT sites *in situ* under near-native conditions with the aim of resolving how HIV-1 viral particles efficiently transfer between infected and healthy T cells (Figure 2). By using cryoET, sub-nanometer resolutions can be reached *in situ* while preserving viral particle and cellular ultrastructure. NL43 IRES-eGFP infected SupT1 R5 donor cells (GFP+) were co-cultured with CTDR-stained target SupT1 R5 cells (CTDR+) at a ratio of 2:1. At 3 hours post-mixing, the cells were fixed with 4% PFA, applied to EM grids, and vitrified via plunge freezing. We identified cell-cell interfaces between infected and target cells using cryo fluorescent light microscopy for downstream image registration (cryoCLEM, Figure 2B). Cell-cell interface regions were then processed via cryo focused ion beam (cryoFIB) milling to generate ∼200 nm thick lamellae (Figure 2C and D). Following cryo transmission electron microscopy screening of the lamellae for the presence of particles at the cell-cell interface (cryoTEM, Figure 2E and F), we performed cryoelectron tomography (cryoET) to collect tilt series, which were subsequently reconstructed into cryotomograms. For this study, we defined a CCT site as an intercellular space that contains at least one viral particle in relative proximity to the infected and target cell membranes.

**Figure 2.**
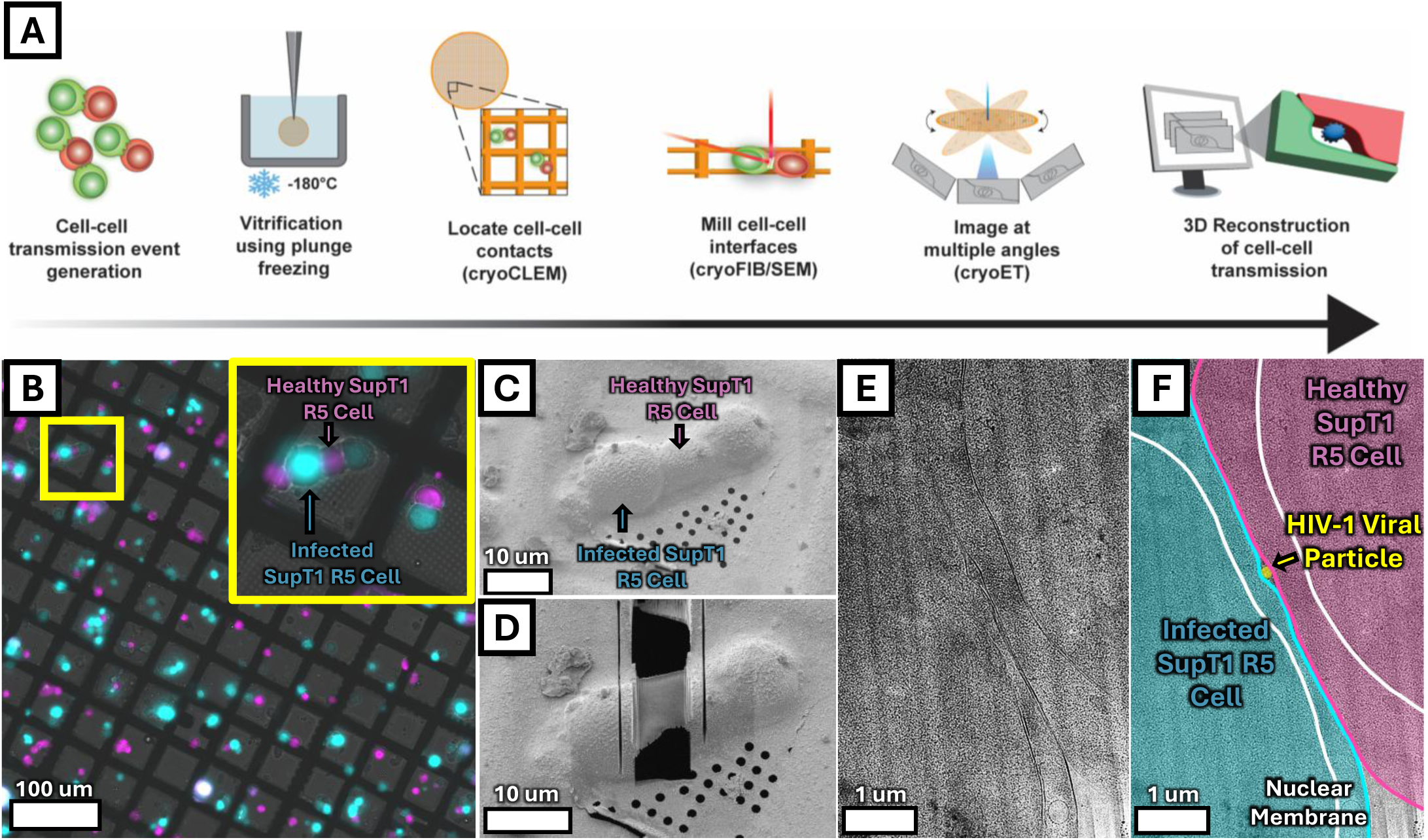
*In situ* correlative cryoET workflow for imaging HIV-1 CCT sites. (**A**) CCT events were generated by co-culturing NL4-3 IRES-eGFP infected donor and uninfected CTDR+ target SupT1 R5 cells for 3 hours. Cells were deposited on to gold electron microscopy grids and vitrified via plunge freezing. GFP-CTDR cell pairs were identified by cryofluorescence microscopy (cryoCLEM), transferred to a cryoFIB/SEM microscope, and milled to generate a thin lamella of the cell-cell interface. The grid was then transferred for overview screening using cryoTEM and tilt-series acquisition using cryoET followed by tomogram reconstruction. (**B**) Cryofluorescence microscopy to identify NL4-3 IRES-eGFP infected donor cells (cyan) and uninfected targets (magenta). Inset highlights a potential CCT conjugate centered within a grid square. (**C, D**) Scanning electron micrographs show the target region from the inset in (B) during cryoFIB/SEM site preparation, captured before (**C**) and after (**D**) focused ion beam milling. (**E, F**) Medium-magnification cryoTEM overviews of the resulting lamella in both raw (**E**) and annotated (**F**) formats.

Of the 20 lamellae assessed, 10 lamellae were confirmed to have evidence of at least one viral particle in the intercellular space between the target and infected SupT1 R5 cell (Figure 3A and B). From these lamellae, 16 tomograms were collected total that were considered CCT sites. On average, 2.4 ± 1.3 viral particles were found in these sites in either the immature, mature, or budding assembly state (Figure 3C and D). Interestingly, intercellular vesicles were also sometimes found in these spaces, 0.63 ± 0.78 per site (Supplemental Figure 1). Of note, all open intercellular spaces observed contained at least one viral particle or intercellular vesicle. This absence of empty open spaces suggests that HIV-1 CCT sites may require the presence of at least one intercellular particle to be supported, whether that be a viral particle or vesicle.

**Figure 3.**
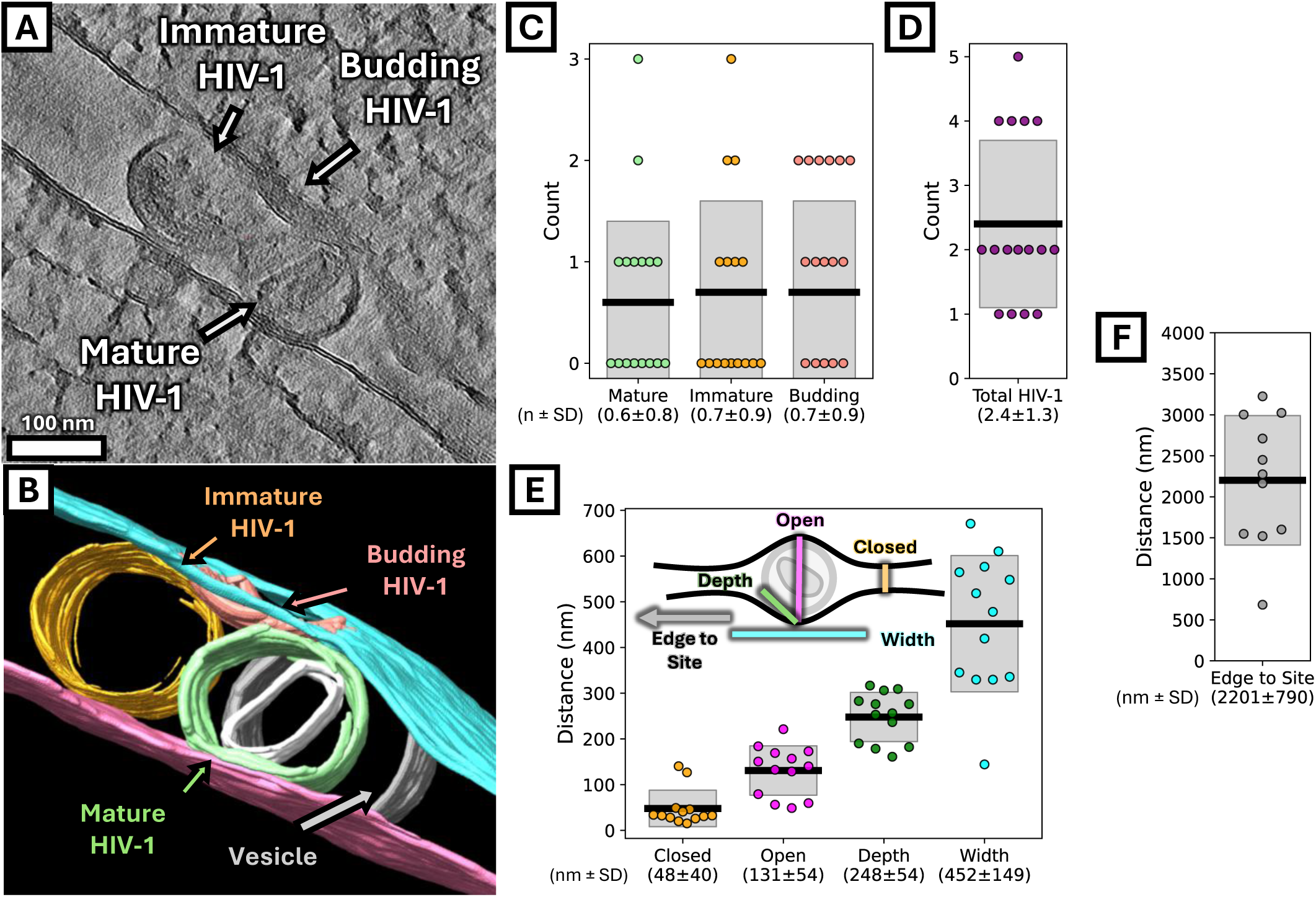
HIV CCT sites are small and contain few viruses. (**A, B**) A representative tomographic slice (**A**) and corresponding three-dimensional segmentation (**B**) displaying a CCT site containing budding, immature, and mature HIV-1 particles and an intercellular vesicle. (**C**) Quantification and distribution of HIV-1 across different maturation and assembly stages within the collected tomograms. Each dot represents one tomogram. (**D**) The total numbers of viral particles (including budding viral particles) per CCT site. (**E**) Dimensions of CCT site as shown in inset. (**F**) The distance from the CCT site to the remaining extracellular space past the cell edge. Values below x-axis indicate the mean and standard deviation for each category. Black bar represents the mean and the gray box represents ±1 standard deviation.

To further characterize the ultrastructure of these HIV-1 CCT sites, we measured several dimensions of each 3D reconstruction and their relative position in their respective cell-cell interface (Figure 3E) in the lamella overview. The distance between healthy and infected cell membranes outside the virus-containing volume (Closed) was measured to be on average 48 ± 40 nm while the distance between membranes within the virus-containing volume (Open) was on average 131 ± 54 nm. The distance between the bounds of the virus containing volume (Width) as well as the depth of each volume (Depth) was measured to be 452 ± 149 nm and 248 ± 54 nm, respectively. When looking at cryoTEM overview images of confirmed CCT sites, we found that these small sites are largely secluded from the remaining extracellular space being on average 2.2 ± 0.79 μm away from the cell-cell interface edge (Figure 3F).

Previous studies have utilized resin-embedded transmission electron microscopy (reTEM) to better characterize the CCT interfaces (4, 58). To ensure that the differences in cell morphology were due to our cryoET sample preparation and not our CCT model system, we imaged CCT events using reTEM (Supplemental Figure 2). Due to the lack of fluorescence after resin-embedding, we relied on the visual observation of HIV-1 particles and budding profiles to help identify the infected SupT1 R5 cells. In the reTEM micrographs, we observed more membrane ruffling, more budding profiles, and increased distance between infected and target cell membranes as compared to cryoET prepared samples. Morphological differences in the membranes also extend to the nuclear envelope. This difference in morphology between cryoET and reTEM suggests that the imaging preparation methodology selected directly impacts the observations at the cell-cell interface, and therefore, at the CCT sites themselves.

In our cryoET observations, we observe viral particles at multiple stages of assembly, budding and maturation. While each particle is an independent virus, we can use the completion stage of the Gag lattice to fit together a model for CCT site biogenesis. Our data indicates that the assembly of HIV-1 particles at the cell-cell interface may be a major driver for CCT site generation (Figure 4, Supplemental Movie 1). During viral assembly, the infected cell membrane is deformed through lateral Gag-Gag interactions necessary to build the immature lattice. To accommodate the curvature of the immature lattice, the infected cell membrane is pulled away from the target cell membrane, creating the small open space for CCT with the assembling virus in its center. Following successful budding, the virus is released from the infected cell membrane and the CCT site is complete. Other viral particles may also assemble and bud at this site. In the CCT sites, HIV-1 undergoes maturation, converting from the immature to the mature state. Altogether, our data shows that CCT sites imaged under near-native conditions are small, contain very few viral particles, and are separated from the extracellular space by long, straight and narrow interfaces between the infected donor and uninfected target cells.

**Figure 4.**
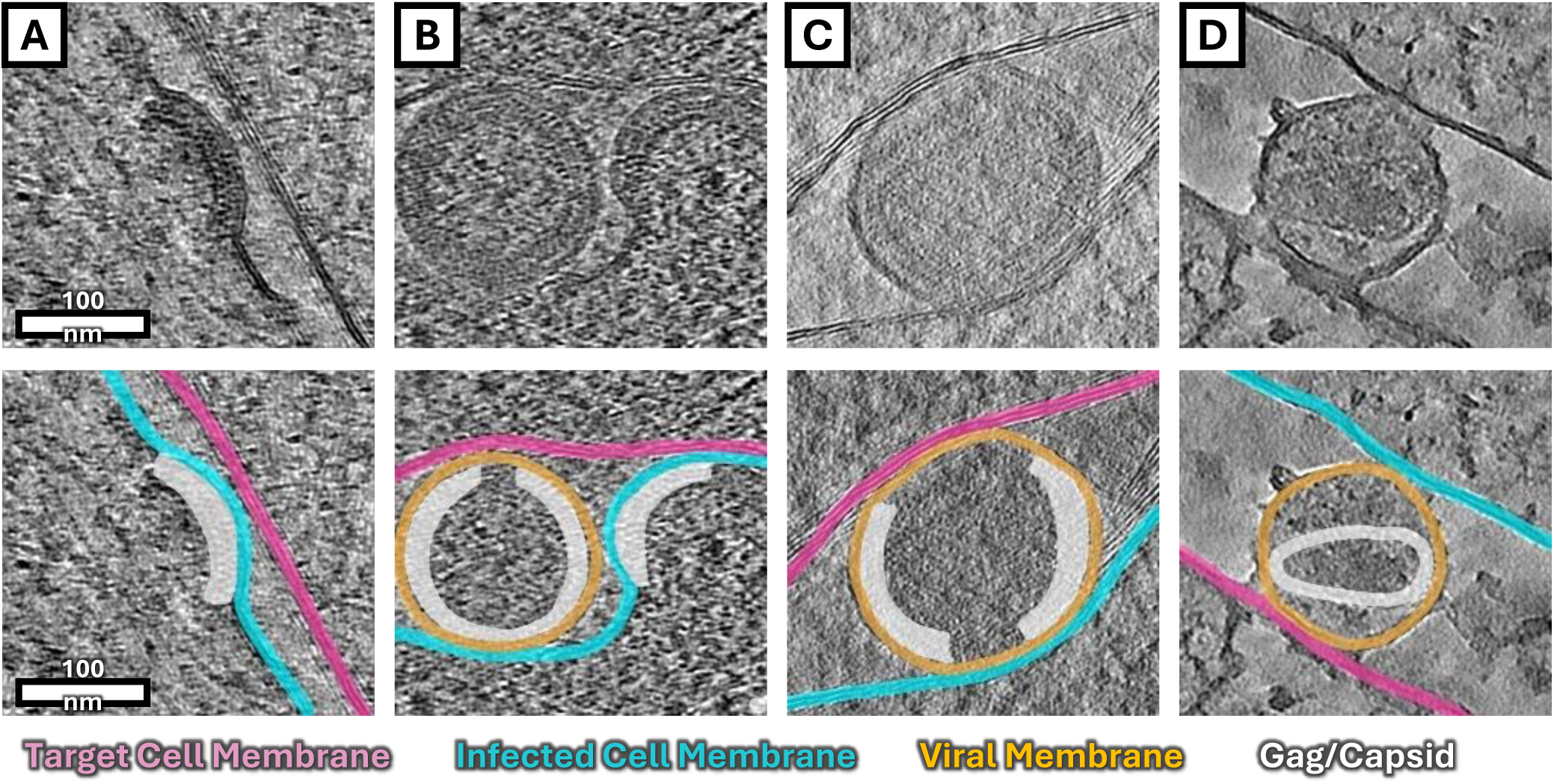
Tomographic slices depicting the progression of viral assembly within CCT sites. (**A–D**) Tomographic slices capturing sequential stages of HIV-1 assembly, including early assembly (**A**), deformation of the infected cell membrane during advanced assembly (**B**), a fully budded immature virus (**C**), and a mature virus (**D**). The corresponding bottom panels display the annotated versions of each top panel for highlighting specific structural features.

### Multicell conjugates are the major GFP+CTDR+ population at 3 hours but not 24 hours

One of the major determinants of CCT is the presence of HIV-1 Envelope on the infected cell surface. To complement our imaging workflow, we wanted to further validate that cell-cell interfaces generated at 3 hours post mixing were dependent on HIV-1 Envelope and not a byproduct of background T cell interactions. For this, we generated the Env-deficient mutant ΔEnv IRES-eGFP which removed a majority of gp120 and frameshifted the remaining Env sequence to produce a premature stop codon (Figure 5A). As this deletion causes the produced viruses to be non-infectious, we included a VSV-G envelope during viral particle production. This allowed us to generate a single cycle infectious donor population prior to mixing with CTDR+ target cells.

**Figure 5.**
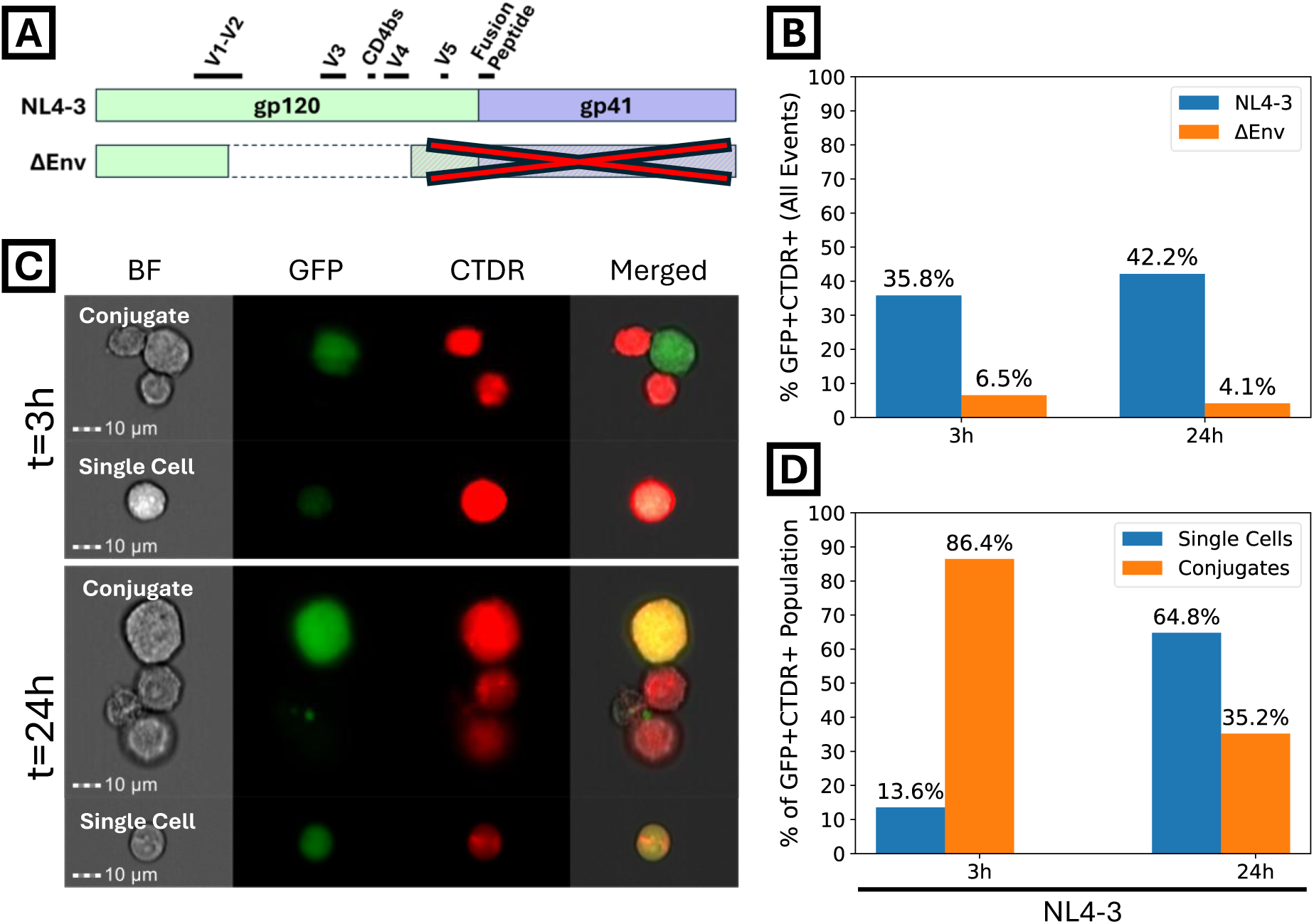
Population composition of HIV-1 CCT co-cultures analyzed by imaging cytometry. (**A**) Comparison NL4-3 Envelope and **Δ**Env sequences. Dotted box indicates sequence deletion and hashed box with red cross indicates lack of translation due to frameshift and introduction of a stop codon (**B**) The percentage of all (single-cell and conjugate) GFP+CTDR+ events at 3h or 24h co-cultures of uninfected SupT1 R5 cells mixed with either NL4-3 IRES-eGFP or **Δ**Env IRES-eGFP infected SupT1 R5 cells. (**C**) Representative imaging cytometry of single cells and conjugates present in NL4-3 IRES-eGFP and **Δ**Env IRES-eGFP co-cultures at 3h and 24h. (**D**) Breakdown of the distribution of single cell and conjugate GFP+CTDR+ events within the 3-hour or 24-hour co-cultures. Data from N=1 experiment that was performed in duplicate.

To assess CCT in a quantitative manner, we sought to develop a flow cytometry assay which would allow for relative quantification of CCT events at the population level. As flow cytometry is more traditionally used to assess single cells instead of multicell conjugates, we first wanted to be sure that CCT events could persist during flow cytometry. To address this and to better understand the co-culture subpopulations, we generated co-cultures containing NL4-3 IRES-eGFP and **Δ**Env IRES-eGFP (similar to our cryoET samples) and used the Amnis ImageStream imaging cytometry system to image events after 3 and 24 hours. By using both timepoints, we can assess at 3 hours if CCT conjugates survive flow cytometry and at 24 hours if newly infected cells were present. At 3 and 24 hours, NL4-3 IRES-eGFP had 29.3% and 38.1% more GFP+CTDR+ events as compared to **Δ**Env IRES-eGFP, respectively (Figure 5B). This increase in the number of GFP+CTDR+ events due to the presence of HIV-1 Envelope points to our co-culturing system being able to produce Env-dependent events and therefore most likely HIV-1 CCT events.

To further characterize the GFP+CTDR+ populations at 3 and 24 hours, we assessed images for the presence of multicell conjugates and single cells at each timepoint (Figure 5C and D). At 3 hours, we saw that 86.4% of GFP+CTDR+ events were a multicell conjugate containing at least one target CTDR+ and one infected GFP+ cell. In contrast, at 24 hours, newly infected single cells make up 64.8% of double positive events. Importantly, in the 35.2% of events that were considered multicell conjugates at 24 hours, only 28.75% were between a strictly GFP+ and CTDR+ cell. This indicates that the majority of multicell conjugates at 24h are not conjugates of the initially infected population interacting with target CTDR+ cells but instead other infection-related outcomes including newly infected cells forming conjugates or syncytia-related events (Supplemental Figure 3). These results suggest that CCT conjugates are able to endure flow cytometry analysis and that by 24 hours, co-cultures are able to produce successful infection events.

### CCT conjugates show Env-dependence in primary CD4+ T cells using conventional flow cytometry

With validation of CCT events at 3 and 24 hours with imaging cytometry, we next wanted to use conventional flow cytometry with NL4-3 IRES-eGFP and **Δ**Env IRES-eGFP to increase our throughput and confirm the Envelope dependence in SupT1 R5 and primary CD4+ T lymphocytes. Based on our imaging cytometry results, for 3-hour analysis we focused only on analyzing multicell conjugates (CCT conjugate formation) as these were the majority of double positive events at 3 hours. Due to the heterogeneity of the 24-hour population of GFP+CTDR+ non-single cell events with imaging cytometry, we decided to assess infection efficiency by focusing only on GFP+CTDR+ single cells at 24 hours, similar to previous work (59). To distinguish single cells from multicell conjugates, we utilized a gating strategy using forward and side scatter with an emphasis on signal width as suggested by previous work (60) (Supplemental Figure 4). After single and multicell conjugate separation, each subpopulation was then assessed for GFP and CTDR fluorescence with the population of interest being events that were considered both GFP+ and CTDR+ (Figure 6A, C, E and G). Initial co-culture conditions at 0 hours were assessed using single cell and multicell conjugate gating as a point of comparison for the 3-hour and 24-hour co-cultures (Supplementary Figure 5).

**Figure 6.**
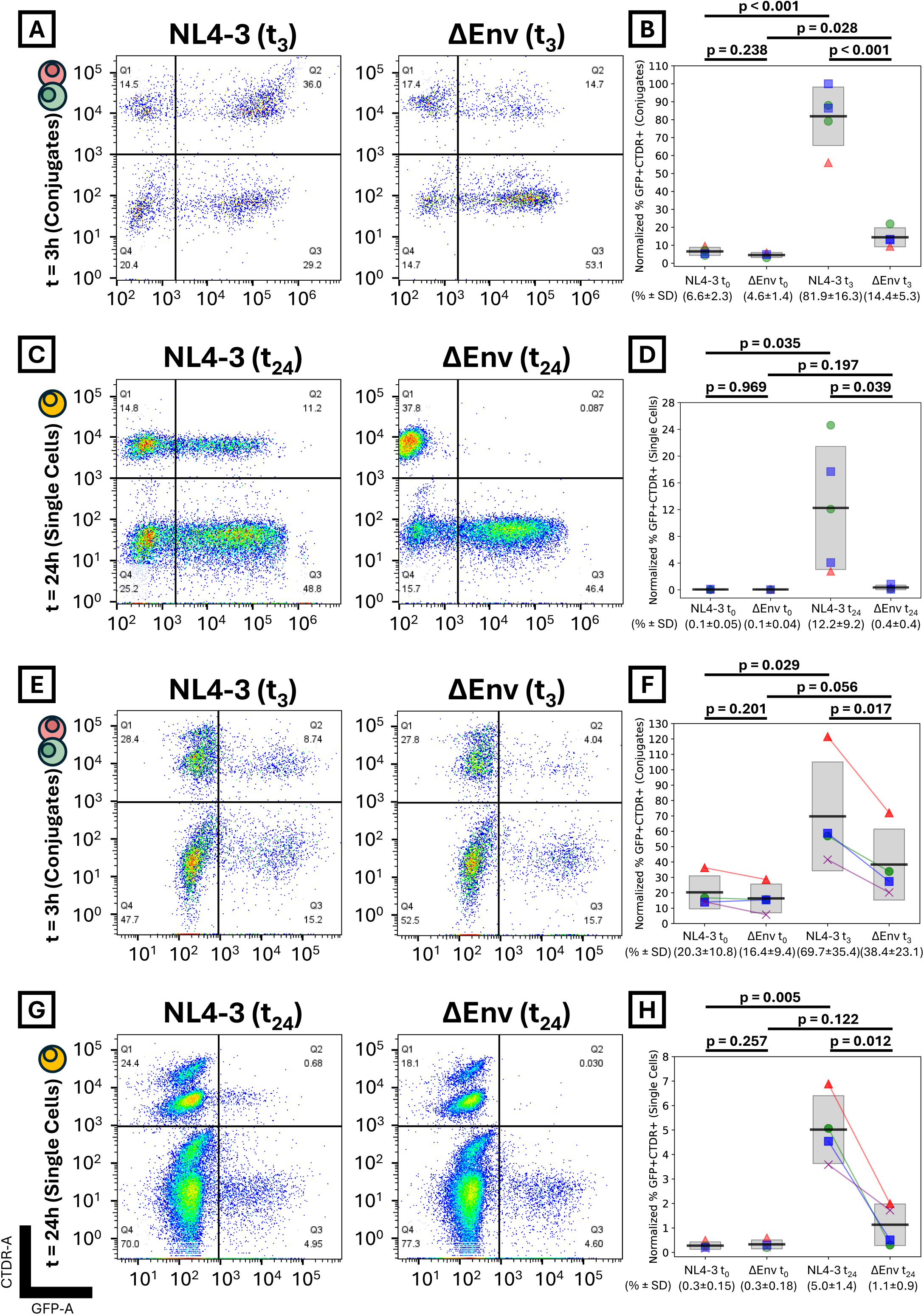
Comparison of NL4-3 and ΔEnv CCT using conventional flow cytometry. (**A**) Representative dot plots of multicell conjugates in 3h co-cultures between target CTDR+ SupT1 R5 cells mixed with either NL4-3 IRES-eGFP or ΔEnv IRES-eGFP infected SupT1 R5 cells. (**B**) The percentage of GFP+CTDR+ multicell conjugate events from initial 0-hour or 3-hour co-cultures. (**C**) Representative dot plots of single cells in 24-hour co-cultures of target CTDR+ SupT1 R5 cells mixed with either NL4-3 IRES-eGFP or ΔEnv IRES-eGFP+ infected SupT1 R5 cells (**D**) The percentage of GFP+CTDR+ single cell events from initial 0-hour or 24-hour co-cultures. Distinct shapes in **B** and **D** each represent distinct biological replicates (N = 3) and each individual shape a technical replicate. **(E-H)** Same as **A-D** but with co-culture of infected GFP+ primary CD4+ T lymphocytes and target CTDR+ primary CD4+ T lymphocytes. Shapes in **F** and **H** represent distinct primary cell donors (N = 4). All values are normalized to initial GFP+ infection levels prior to co-culture. Values below x-axis indicate the mean and standard deviation for each category. Black bar represents the mean and the gray box represents ±1 standard deviation. P-values calculated using either two-tailed unpaired *t-*test (**B** and **D**) or two-tailed paired *t-*test (**F** and **H**).

For both SupT1 R5 and primary CD4+ T lymphocytes, at 0 hours there was no significant change in the number of double positive multicell conjugates (Figure 6B and F) or double positive single cells (Figure 6D and H) when comparing NL4-3 IRES-eGFP and **Δ**Env IRES-eGFP. At 3 hours, NL4-3 IRES-eGFP had a significant increase in the number of double positive multicell conjugates in both SupT1 R5 (Figure 6B, p<0.001) and primary CD4+ T lymphocytes (Figure 6F, p=0.017) when compared to **Δ**Env at 3 hours. Comparing the initial co-culture conditions at 0 hours to those at 3 hours, there was an increase in the number of double positive conjugates in both SupT1 R5 and primary CD4+ T lymphocytes for both NL4-3 IRES-eGFP and **Δ**Env IRES-eGFP. This increase in interactions may be explained by the usual interactions commonly exhibited by T cells but is evident that interactions were markedly higher in Env-competent viruses.

The absence of a fully functional Env led to a significant reduction of infection at 24 hours for SupT1 R5 (Figure 6D, p=0.039) and primary CD4+ T lymphocytes (Figure 6H, p=0.012), as expected, and confirming that our conventional cytometry assays can detect and measure the contribution of molecular determinants of CCT. There were minimal double positive single cells for both viruses present in their initial co-culture conditions (Supplementary Figure 5). Overall, the 3-hour and 24-hour observations with flow cytometry indicate that with HIV-1 Env present there is a significant increase in the number of conjugates and successful infection events. As CCT events are known to be driven by the presence of Env, we concluded that our methodology can successfully quantify CCT events. Because we use equivalent co-culturing methods for cryoET, these findings have also increased our confidence that the infected GFP+ and target CTDR+ multicell conjugates in cryoET are in fact bona-fide CCT events. Moreover, due to the requirement of cell incubation to form a significant number of multicell double positive conjugates, we also can confirm that GFP+CTDR+ events generated at 3 hours and 24 hours were not a byproduct of crosslinking due to paraformaldehyde fixation.

### CCT conjugate formation is unequally affected by Env-CD4 blockade

To further refine our investigation of the Env-CD4 dependence of HIV-1 CCT co-cultures, we treated initial NL4-3 IRES-eGFP infected SupT1 R5 co-cultures with either soluble CD4 (sCD4) to engage HIV-1 Env or anti-CD4 monoclonal antibody SIM.2 to target cellular CD4 (Figure 7). We included a free-virus transmission (FVT) alongside 24-hour co-cultures to ensure and measure the inhibitory effects of each treatment. Using our flow cytometry strategy, sCD4 had no significant impact on CCT conjugate formation (Figure 7A) but dose-dependently impeded productive new CCT infections at 24 hours (Figure 7B). A reduction in infection was also seen in FVT after 24 hours (Figure 7C). Next, we treated co-cultures with SIM.2 to inhibit CD4 binding of HIV-1 Env. Upon increasing concentrations of SIM.2, we observed a significant decrease in newly infected cells in 24-hour co-cultures (Figure 7E) as well as FVT (Figure 7F). For 3-hour co-cultures, CCT conjugates were partially, but not completely, abolished. Our results show that blockade of either Env or CD4 significantly decreases infection in both 24-hour co-cultures and FVT. Interestingly, CCT conjugate formation resists Env blockade with soluble CD4 and partially resists CD4 blockade with SIM.2, pointing towards an asymmetry between Env-CD4 interactions in a cell-cell and in a virus-cell context.

**Figure 7.**
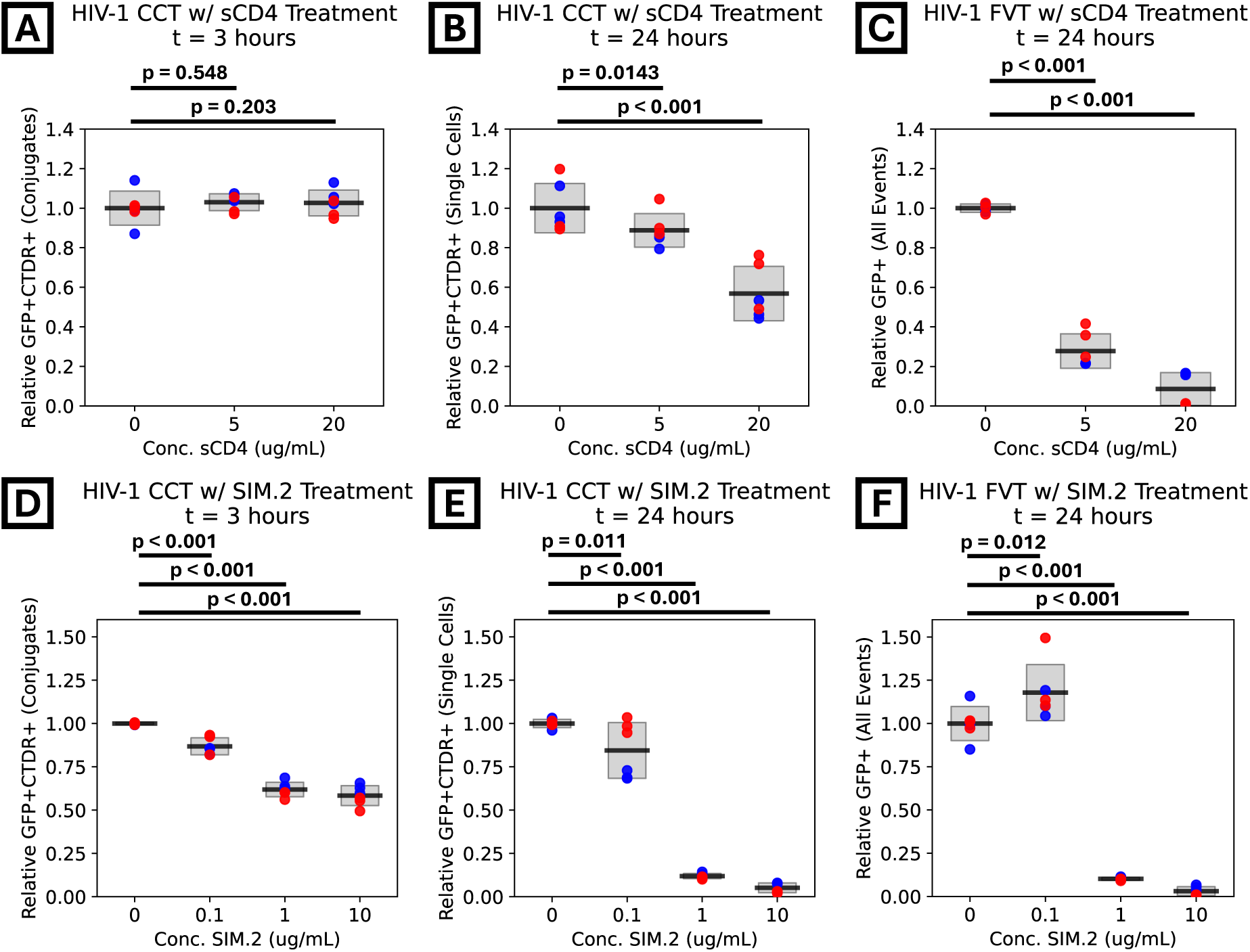
Soluble CD4 and anti-CD4 antibody impact in HIV-1 CCT co-cultures. (**A**, **D**) Quantification of GFP+CTDR+ multicell conjugates events in 3-hour co-cultures of target CTDR+ SupT1 R5 and NL4-3 IRES-eGFP infected SupT1 R5 cells treated with varying concentrations of soluble CD4 (sCD4, **A**) or anti-CD4 (SIM.2, **D**). (**B** and **E**) Quantification of GFP+CTDR+ single cells in 24-hour co-cultures treated with varying concentrations of sCD4 (**B**) or SIM.2 (**E**). (**C** and **F**) Quantification of GFP+ SupT1 R5 after cell-free NL4-3 IRES-eGFP virus infection treated with varying concentrations of sCD4 (**C**) or SIM.2 (**F**). Data are derived from distinct biological replicates (N = 2), where colors represent distinct biological and each dot a technical replicate. All values are normalized to untreated controls and initial GFP+ infection levels prior to co-culture (if applicable). Values below x-axis indicate the mean and standard deviation for each category. Black bar represents the mean and the gray box represents ±1 standard deviation. P-values calculated using one-way ANOVA with a Tukey’s test for multiple comparisons.

## Discussion

Here we provide an ultrastructural analysis of HIV-1 CCT and quantified these interactions using flow cytometry. Using a correlative, near-native, *in situ* imaging approach with cryoET, we showed that HIV-1 CCT sites are small secluded intercellular spaces that contain few viral particles. With our flow cytometry methodology, we decoupled CCT conjugate formation from CCT infection and observed that Env is essential for both multicell conjugate formation and productive HIV-1 infection. However, the interactions between HIV-1 Env and cellular CD4 during CCT conjugate formation responded differently to blockade as compared to FVT. Together, our ultrastructural and cytometry data reveal how the physical architecture of CCT sites and the molecular interactions that establish them jointly govern the efficiency and resistance of cell-cell transmission.

Our cryoimaging observations of CCT sites differ from previous descriptions obtained using conventional resin-embedded electron microscopy. Prior studies using resin embedding have described spacious intercellular compartments containing dozens of viral particles with scalloped and interdigitated membranes between infected and target cells (4, 31, 33, 58). Although viral particles occupy these compartments, large regions of apparent “empty” space were also typically observed. In contrast, our cryoimaging data consistently revealed highly confined interfaces containing at most 5 viral particles per site. These compartments were only large enough to accommodate the particles themselves which were commonly in direct contact with both opposing cellular membranes and occasionally visibly compressed between the membranes. Viral particles showed no detectable void space around them unless a larger vesicle was supporting the site. Outside of the virus-containing regions, the membranes of infected and target cells remained in very intimate contact, separated by less than 50 nm over an extended distance often exceeding 2 µm before opening into the remaining extracellular space. Unlike the scalloped morphology reported previously, these membranes appeared smooth and relatively straight. We believe that these differences were largely attributable to the distinct sample preparation procedures used for cryoimaging versus conventional resin-embedding electron microscopy. Consistent with this interpretation, when our samples were processed using traditional resin-embedding methods, we observed features resembling those reported previously, including scalloped membranes and enlarged intercellular compartments. Resin-embedding requires dehydration and resin curing steps that can induce substantial sample shrinkage, up to 60% (61). Such an effect could disrupt the close membrane-membrane interfaces observed under hydrated cryogenic conditions by physically pulling opposing membranes apart during sample shrinkage, while also potentially merging adjacent contact sites into larger, more spacious compartments, containing a large number of viral particles. In addition, most conventional resin-embedding abolishes fluorescence signals, complicating identification of infected cells. As a result, site selection necessarily relied on visual screening for prominent Gag lattices in infected cells and accumulations of viral particles in intercellular spaces. This may bias detection toward larger, virus-rich compartments while overlooking more compact sites containing fewer viral particles. We therefore suggest that prior descriptions of spacious, virus-rich compartments reflect preparative artifacts rather than the native architecture of cell-cell transmission sites. Taken together, our findings highlight how cryoimaging can reveal the native organization of cell–cell transmission sites under near-physiological, hydrated conditions, preserving membrane morphology and intercellular spacing that are likely sensitive to preparative artifacts.

Unfortunately, cryoimaging captures only a thin lamella (∼200 nm thick) from a larger three-dimensional cellular interface, such that the full extent and organization of individual CCT sites cannot be visualized in a single tomogram. Based on the average thickness of our lamellae, the likelihood of finding a CCT site in our lamellae, the average length of cell-cell contact between an infected and an uninfected cell and the average number of viruses found in a CCT site, we estimate that dozens of viruses are transmitted in the entire cell-cell contact. This is still lower than numbers estimated from prior fluorescence microscopy and resin-embedding TEM studies (5, 12, 16, 23, 27–31). Another potential limitation is that the inherently low signal-to-noise ratio of *in situ* cryoET data can obscure more subtle structural features within the tomograms. The demanding sample preparation workflow and low throughput of cryoimaging further restricts the number of sites that can be analyzed, making our observations primarily qualitative rather than quantitative. Despite these limitations, cryoimaging currently remains the only approach capable of three-dimensionally visualizing these interfaces within fully hydrated, near-native cellular environments at nanometer-scale resolution.

Based on this native architecture, we propose a model where HIV-1 CCT efficiency and broad protection stems from the ultrastructure of the CCT intercellular space. Viral particles assemble, bud, and mature in these confined spaces in close proximity to the target cell membrane, positioning virions for immediate engagement with the target cell membrane receptors. Of note, we did not observe viral entry intermediates in our dataset, consistent with the known rapid kinetics of HIV-1 entry and the general challenge of capturing these transient events by cryoET across different viral systems (62–66). If virion transfer across the confined interface is rapid, capturing these entry events via cryoET is inherently unlikely. Furthermore, the efficiency of CCT may directly reflect the kinetic advantage provided by this architecture. Indeed, live imaging studies of HIV-1 CCT show rapid transfer of Gag across CCT interfaces, consistent with our absence of observed entry intermediates (67). Similarly, the same structural confinement that positions virions for rapid entry may also shield them from neutralizing antibodies and antiviral agents. The secluded virus-containing sites surrounded by narrow and long intercellular interfaces could physically restrict access of molecules to the viruses, providing a structural basis for the broad protection associated with cell-cell transmission. Direct experimental testing of these hypotheses will be the focus of subsequent studies.

Our cytometry data demonstrated that CCT conjugate formation and infection efficiency are both dependent on Env and CD4 presence, as consistent with prior reports (2, 34). Specifically, deletion of functional Envelope reduced both cell-cell conjugate formation and infection efficiency. Prior cytometry approaches have independently used fluorescent Gag transfer assays and molecular clones expressing GFP to investigate transmission steps and productive infection during CCT (32, 59). While these studies primarily focused on reporter protein expression in target cells to quantify infection patterns, the starting point of CCT (i.e. initial cell-cell interface formation) was not investigated in a quantitative manner. By including non-single cell populations, our cytometry workflow advanced on these approaches by parsing out the importance of Env interactions in both initial conjugate formation as well as productive infection within a single assay. This is a critical consideration given that CCT is fundamentally a reliant process on interacting cells.

We also demonstrated that blocking viral gp120 with soluble CD4 and blocking the native CD4 receptor with SIM.2 antibody has distinct effects on CCT versus FVT, with sCD4 inhibiting FVT more potently than CCT at 24 hours. This is consistent with prior observations using antibodies targeting the CD4 binding site in gp120, which similarly show reduced potency against CCT compared to FVT (17). This supports the conclusion that the CCT intercellular interface presents a distinct molecular environment from that encountered during FVT. Most notably, sCD4 and SIM.2 treatments differed in their effects on CCT cluster formation at 3 hours: Env inhibition via sCD4 had no impact, whereas CD4 inhibition via SIM.2 partially reduced conjugate formation. Within the CCT environment, it is likely that Env exists in two distinct pools: cell-bound Env on the donor membrane and virus-bound Env on the virion surface. It is plausible that these two Env populations serve distinct roles: cell-bound Env could predominantly support the intercellular physical contact, while virus-bound Env functions to mediate downstream viral entry. Because sCD4 treatment failed to disrupt conjugate formation, unlike the CD4-targeting SIM.2, it is possible that sCD4 cannot effectively block the cell-associated Env interactions driving this intercellular contact. This functional asymmetry is further reinforced by the results with **Δ**Env genetic knockout where Env was still required for conjugate formation but yet its direct inhibition with sCD4 seemingly had no effect. Overall, our cytometry method can now be applied to systematically interrogate other molecular interactions implicated in CCT, including viral and host factors beyond Env and CD4 (7, 68–70) to determine whether they contribute to the initial cell conjugation step, productive infection, or both. This methodology provides a more granular functional map of the CCT process than has previously been achievable and allows for the relative quantification of CCT stages within the same higher throughput assay.

In summary, this study redefines the physical architecture of HIV-1 cell-cell transmission sites and introduces a quantitative cytometry framework for dissecting the molecular interactions that govern them. The concept of ‘architectural resistance’ emerges from our findings as a necessary consideration for HIV-1 therapeutic development: the narrow, secluded intercellular spaces that characterize CCT sites may act as a barrier to antibody and drug access, providing a structural basis for why certain broadly neutralizing antibodies show reduced efficacy *in vivo* despite high *in vitro* potency against isolated virions. Overcoming this barrier may therefore require evaluating therapeutic candidates not only by their molecular potency, but by their capacity to physically access these restricted cellular interfaces. These structural insights are functionally substantiated by our cytometry workflow, designed to capture the complexity of CCT by decoupling initiation from subsequent transmission and infection. By accounting for multicell events rather than just single cells, our method reflects the dynamics inherent to cell-cell transmission and can be leveraged to revisit other molecular interactions important for CCT with greater accuracy. Collectively, these findings highlight the physical microenvironment as a determinant of intrahost HIV-1 transmission and establish complementary structural and functional tools for interrogating and targeting the cell-cell transmission interface.

