## Supplemental Legends for "Cryoelectron tomography of HIV-1 cell-cell transmission conjugates reveals a secluded environment for viral assembly and transfer"

### Supplemental Figure Legends

**Figure S1. CCT site occupancy including intercellular vesicles.**
(**A**) Quantification and distribution of HIV-1 across different maturation and assembly stages and the number of vesicles within the collected tomograms. Each dot represents an individual CCT site (**B**) The total numbers of viral particles (including budding viral particles) and total number of particles (HIV-1 and intercellular vesicles) per CCT site. Values below x-axis indicate the mean and standard deviation for each category. Black bar represents the mean and the gray box represents ±1 standard deviation.

**Figure S2. Morphology of CCT sites is dependent on sample preparation.**
(**A, B**) Medium-magnification cryoTEM overviews display a CCT site within a lamella milled at the SupT1 R5 cell-cell interface, shown in both raw (**A**) and annotated (**B**) formats. (**C**) A resin-embedded transmission electron microscopy (reTEM) image highlighting the altered membrane morphology of the SupT1 R5 cell–cell interface resulting from conventional dehydration and resin-embedding protocols. Infected SupT1 R5 cells in reTEM were identified using the presence of HIV-1 budding profiles.

**Figure S3. Breakdown of the multicell conjugate population at 24-hour co-cultures.**
(**A**) Representative imaging cytometry panels displaying distinct multi-cell subpopulations within 24-hour co-culture between NL4-3 GFP+ infected and target CTDR+ SupT1 R5 cells. (**B**) Quantification of each identified subpopulation within the 24-hour co-culture. Data displayed is from N=1 experiment performed in duplicate.

**Figure S4. Separation of single and multicell conjugates by conventional flow cytometry.**
For the separation of single and multicell conjugates, events were initially gated for debris removal and for the collection of the majority of events in the FSC-A vs. SSC-A (top left panel), followed by an optimized single cell gate strategy using FSC-W vs. FSC-A, SSC-A vs. SSC-W and FSC-W vs. SSC-W plots. Events that were inside all three gates were considered single cell events (AND Gating) and all other events were considered non-single cell conjugates. Quantification of new infection events was evaluated by the percentage of GFP+CTDR+ single cells out of all single cells at 24 hours. Quantification of CCT conjugation events was evaluated by the percentage of GFP+CTDR+ multicell conjugates out of all multicell events at 3 hours. GFP-A and CTDR-A thresholds were based on initial GFP+ and CTDR+ populations prior to co-culturing. Gating strategy was applied to both SupT1 R5 and primary CD4+ T lymphocytes.

**Figure S5. Paraformaldehyde inactivation does not generate false-positive GFP+CTDR+ conjugates.**
(**A**) Representative flow cytometry plots display the initial co-culture conditions at the 0 hour time point for co-cultures with uninfected and NL4-3 or ΔEnv infected cells in SupT1 R5 and in primary CD4+ T lymphocytes using the gating strategy as described in Figure S4.

**Supplemental Movie 1. Proposed model of HIV-1 CCT site generation.**
Proposed model of HIV-1 CCT site generation where HIV-1 particle assembly deforms the infected cell membrane followed by particle budding and subsequent maturation.
