## Supplementary figures and images for "Cryoelectron tomography of HIV-1 cell-cell transmission conjugates reveals a secluded environment for viral assembly and transfer"

### Supplemental Figures

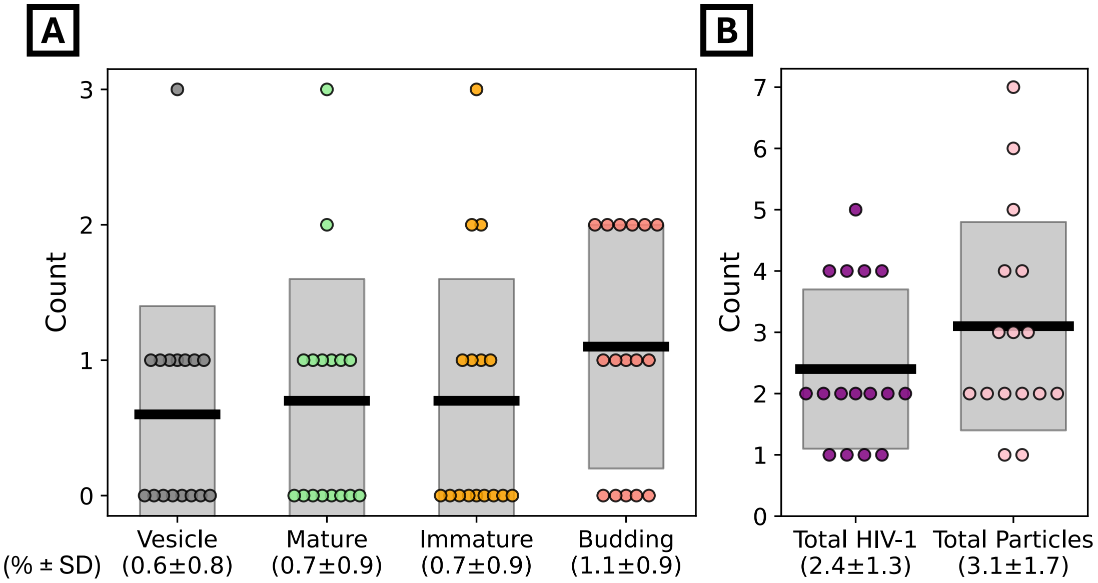

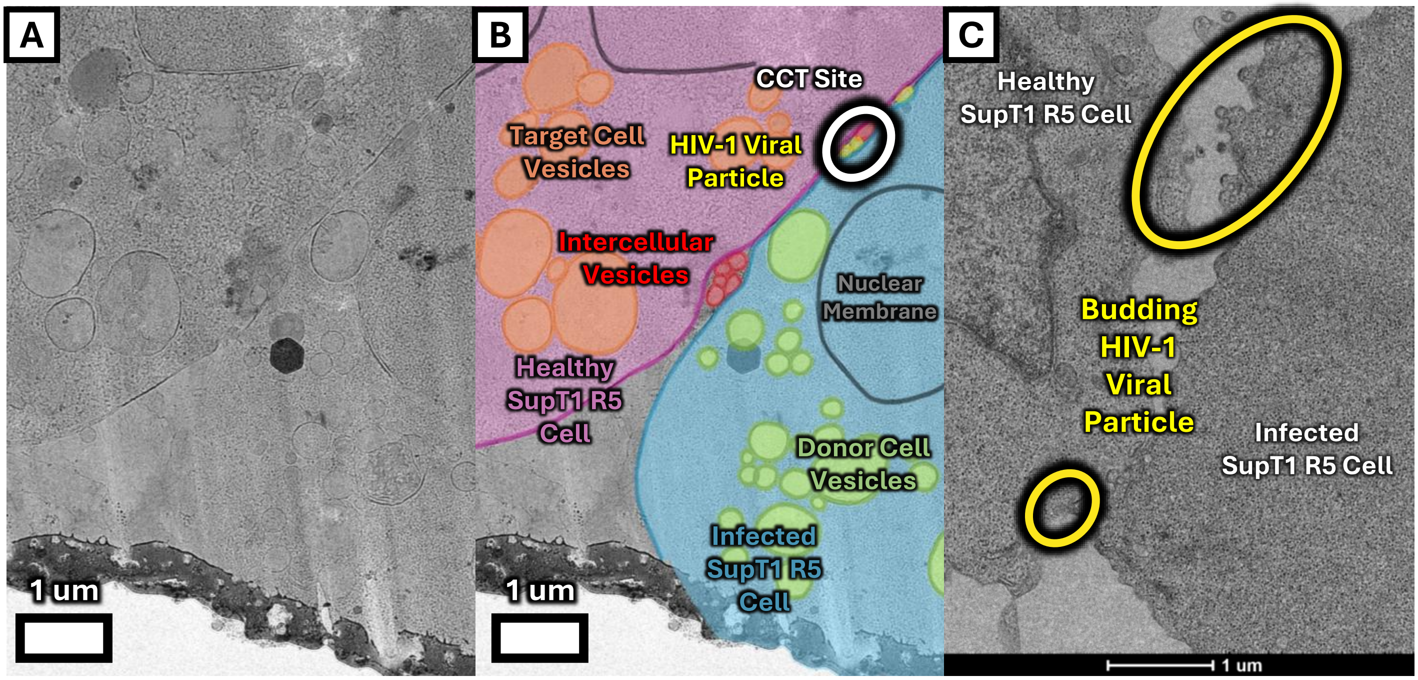

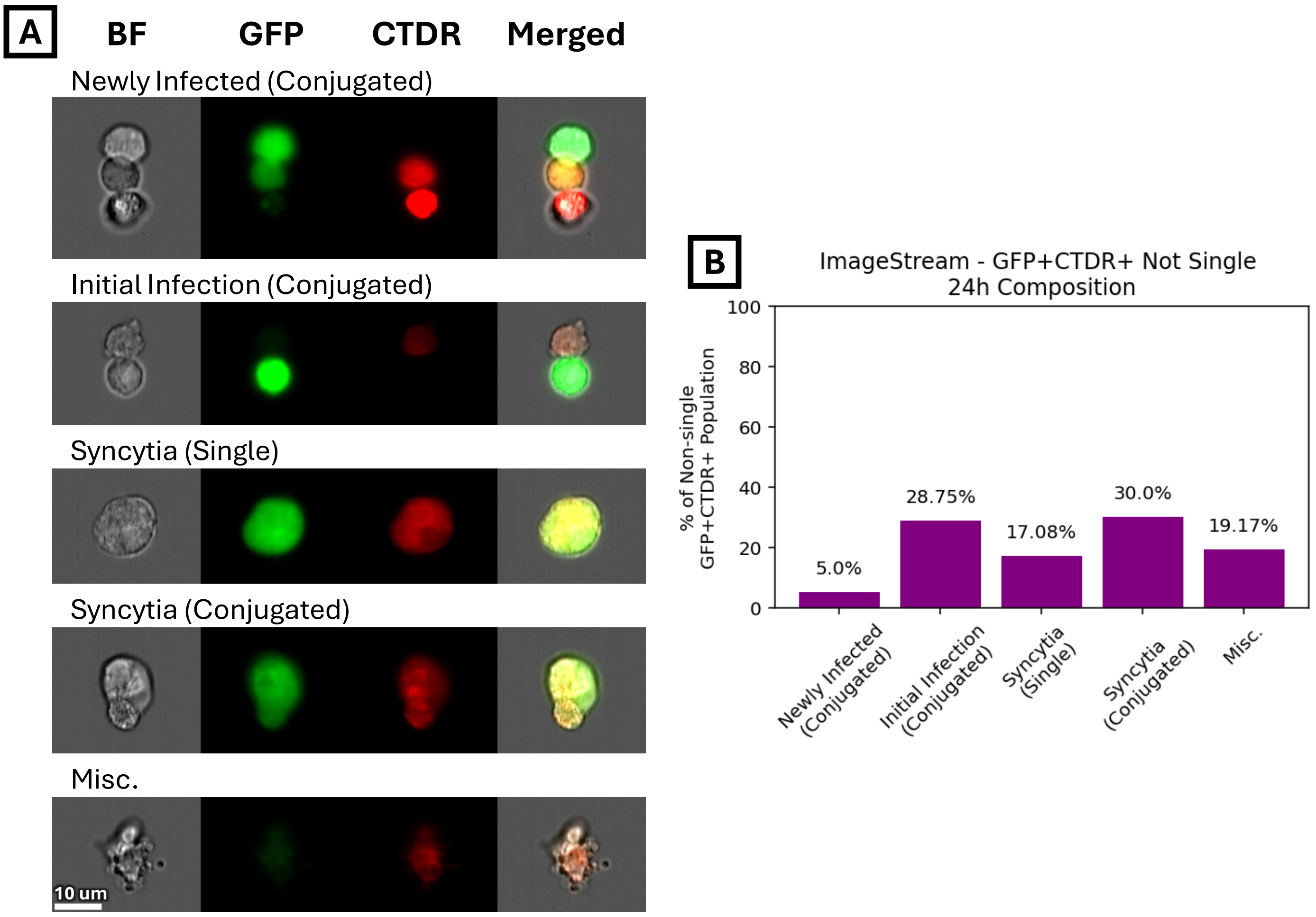

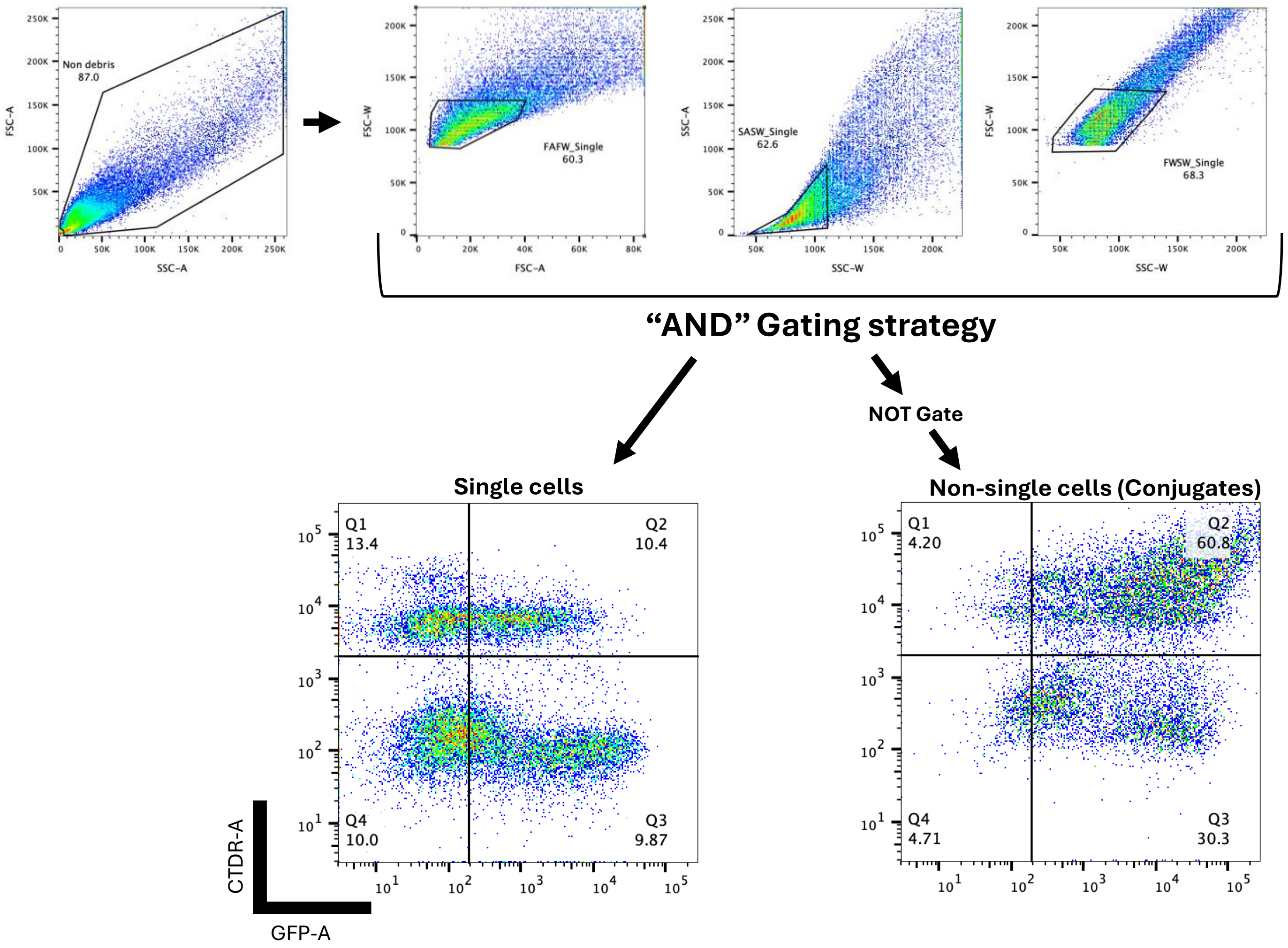

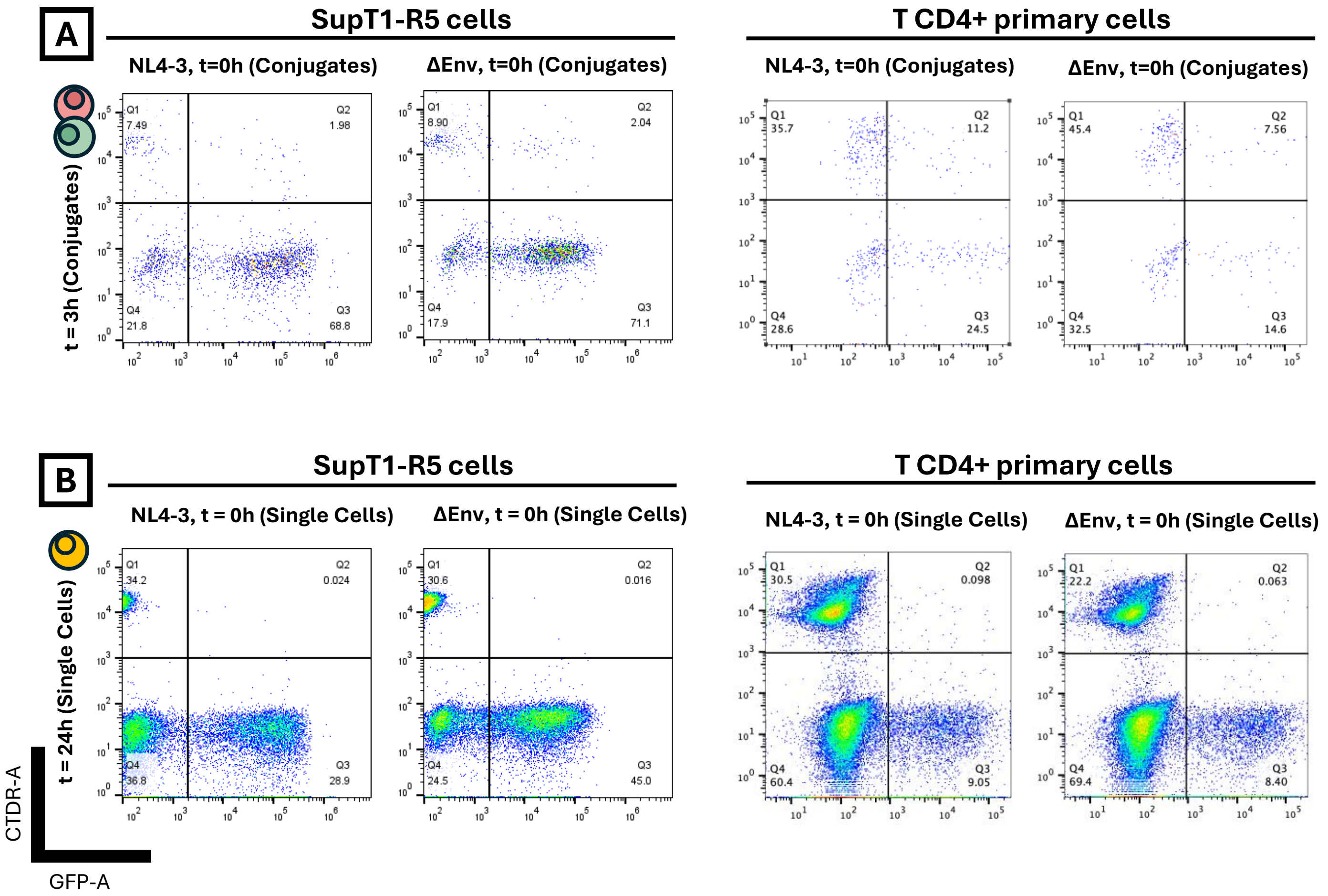
